# Adenine methylation is very scarce in the drosophila genome and not erased by the Ten Eleven Translocation dioxygenase

**DOI:** 10.1101/2023.09.15.557947

**Authors:** Manon Boulet, Guerric Gilbert, Yoan Renaud, Martina Schmidt-Dengler, Emilie Plantié, Romane Bertrand, Xinsheng Nan, Tomasz Jurkowski, Mark Helm, Laurence Vandel, Lucas Waltzer

**Author notes:** co-corresponding authors &. co-first authors (Manon Boulet, Guerric Gilbert, Yoan Renaud).

## Abstract

N6-methyladenine (6mA) DNA modification has recently been described in metazoans, including in drosophila, for which the erasure of this epigenetic mark has been ascribed to the Ten Eleven Translocation (TET) enzyme. Here, we re-evaluated 6mA presence and TET impact on drosophila genome. Using axenic or conventional breeding conditions, we found traces of 6mA by LC-MS/MS and no significant increase in 6mA levels in the absence of TET, suggesting that this modification is present at very low levels in the drosophila genome but not regulated by TET. Consistent with this latter hypothesis, further molecular and genetic analyses showed that TET does not demethylate 6mA but acts essentially in an enzymatic-independent manner. Our results call for further caution concerning the role and regulation of 6mA DNA modification in metazoans and underline the importance of TET non-enzymatic activity for fly development.

## Introduction

Until recently, N^6^-methyl-2’-deoxyadenosine (also called N6-methyladenine or 6mA) was considered to be essentially restricted to the genome of prokaryotes, where this modification plays a well-established role in the restriction-modification system and other processes such as DNA replication or transcription (Sanchez-Romero & Casadesus, 2020; Wion & Casadesus, 2006). Since 2015, several reports detected the presence of 6mA in the DNA of different eukaryotic organisms (Alderman & Xiao, 2019; Boulias & Greer, 2022), including in metazoans (Greer *et al*., 2015; Koziol *et al*., 2016; Liu *et al*., 2016; Wu *et al*., 2016; Xiao *et al*., 2018; Xie *et al*., 2018; Zhang *et al*., 2015a). Although a small fraction of all adenines seems methylated at N6 position (from 0.4% to 0.0001% or below), it was proposed that this modification participates in eukaryotic genome regulation (Wu, 2020). Yet, the significance of 6mA in eukaryotes and the enzymes involved in its metabolism remain controversial with several studies questioning the existence and/or the level of this modification, particularly in metazoans (Douvlataniotis *et al*., 2020; Kong *et al*., 2022; Liu *et al*., 2021; Musheev *et al*., 2020; O’Brown *et al*., 2019; Schiffers *et al*., 2017). Part of the controversy stems from the technologies used to detect 6mA (Boulias & Greer, 2022; Li *et al*., 2021). Notably, antibody-based techniques such as dot blot or DNA immunoprecipitation followed by sequencing (DIP-seq) have been particularly called into question to study low levels of 6mA (Bochtler & Fernandes, 2021; Douvlataniotis et al., 2020; Lentini *et al*., 2018). If liquid chromatography coupled with tandem mass spectrometry (LC-MS/MS) provides a sensitive method to identify 6mA and measure its absolute levels unambiguously, bacterial contaminations can affect the results (Douvlataniotis et al., 2020; Kong et al., 2022; O’Brown et al., 2019). Finally, single-molecule real-time sequencing (SMRT-seq) can detect 6mA presence (and location) on genomic DNA but is prone to give rise to a high false discovery rate when 6mA is rare (Douvlataniotis et al., 2020; O’Brown et al., 2019; Zhu *et al*., 2018).

Notwithstanding, 6mA presence appears strongly supported in drosophila genome (He *et al*., 2019; Ismail *et al*., 2019; Shah *et al*., 2019; Yao *et al*., 2018; Ye *et al*., 2017; Zhang et al., 2015a), where this modification was described to be associated with transposable element silencing and activation of gene transcription (He et al., 2019; Yao et al., 2018; Zhang et al., 2015a). Unexpectedly, 6mA demethylation in drosophila was attributed to the Ten Eleven Translocation (TET) enzyme (Zhang et al., 2015a), a member of the 5-methylcytosine (5mC) dioxygenase family (Iyer *et al*., 2009).

Here, we re-evaluated 6mA levels in drosophila and reassessed the impact of TET on this mark. Using LC-MS/MS, we show that 6mA is present at very low levels in drosophila in axenic conditions and that the absence of TET does not lead to any consistent increase in 6mA levels in the larval central nervous system, nor in the whole larva, the embryo or the adult brain. Furthermore, our genetic and molecular analyses suggest that TET is not involved in 6mA demethylation and that its function during drosophila development is largely catalytic-independent.

## Results and discussion

Previously reported levels of 6mA measured by LC/MS-MS in drosophila ranged from 0.07% to 0.0006% (6mA/A), with the highest levels in the early embryo (Yao et al., 2018; Zhang et al., 2015a). However, a recent study reported much lower levels (0.0002%) even in early embryos and showed that initially reported “high” levels of 6mA were likely due to bacterial contaminations (Kong et al., 2022). Indeed, the contamination of genomic DNA (gDNA) by bacterial DNA is a major confounding factor for LC-MS/MS experiments (Douvlataniotis et al., 2020; Kong et al., 2022; O’Brown et al., 2019). Moreover, the presence of intracellular bacteria can also be a source of 6mA (Douvlataniotis et al., 2020). Along that line, it is worth noting that the genome of *Wolbachia*, a frequent endosymbiont in drosophila, codes for DNA Adenine methyltransferases (Saridaki *et al*., 2011). In addition, 6mA derived from exogenous sources might be incorporated into gDNA *via* the salvage pathway (Musheev et al., 2020), and independently of autonomously-directed adenine methylation (O’Brown et al., 2019; Schiffers et al., 2017). To exclude these possible sources of contamination, we generated germ-free drosophila and reared the larvae on chemically-defined (“holidic”) food devoid of exogenous DNA contribution (see Methods). The absence of exogenous or endosymbiotic bacteria in the resulting flies was confirmed by PCR (Fig. 1a). In these conditions, we observed 0.00025% of 6mA in whole larvae (Fig. 1b) and 0.0005% in the larval central nervous system (CNS) (Fig. 1c). Noteworthy, this corresponds to around 200 to 400 methylated adenines per haplogenome. In addition, similar levels of 6mA were measured in the CNS when non-axenic larvae were reared on classic medium without antibiotic treatment (Fig. 1c), suggesting that contamination by exogenous sources is not a major problem in this tissue. These results confirm the presence of a very small fraction of methylated adenines in drosophila DNA.

**Figure 1.**
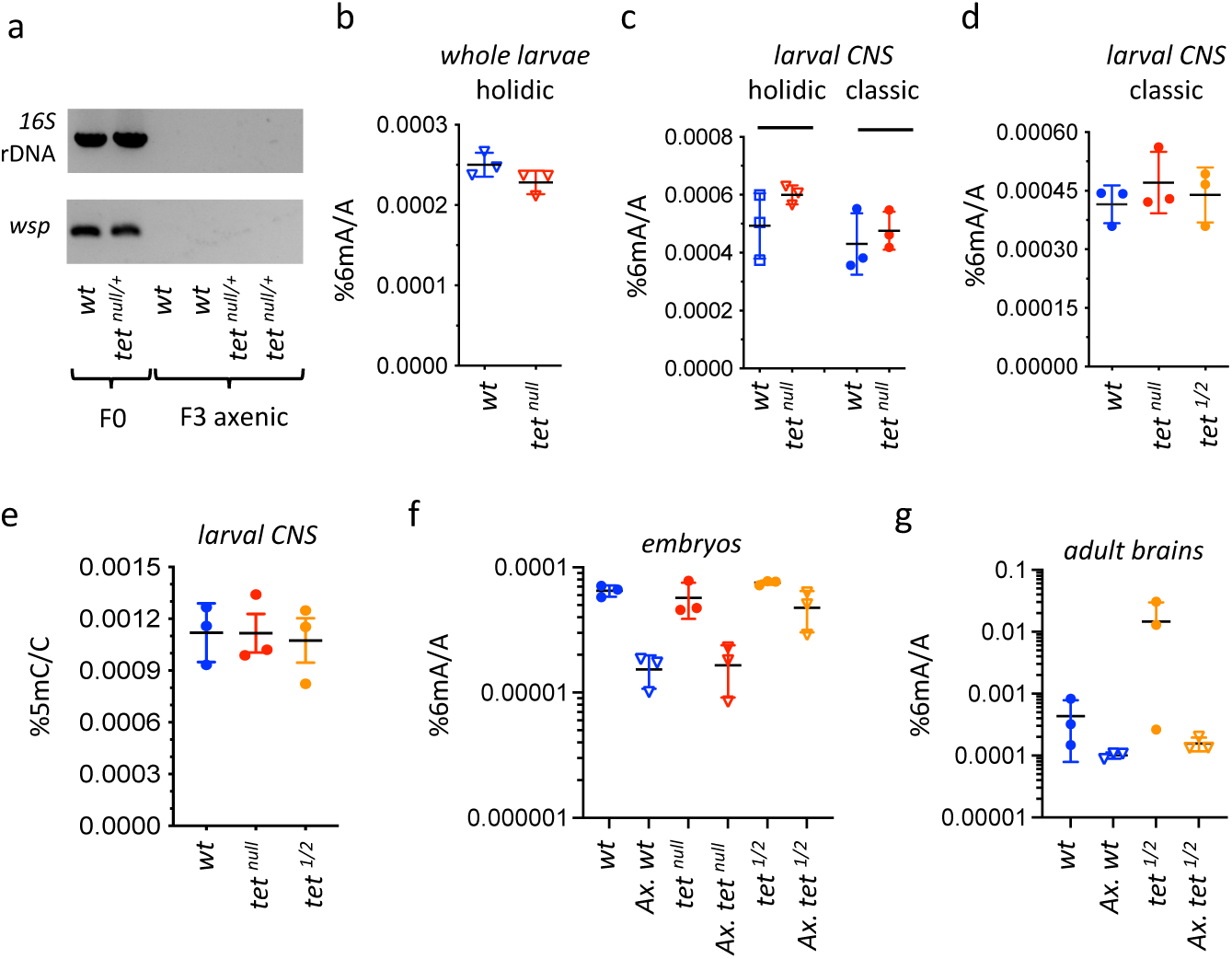
6mA levels in axenic drosophila larvae are very low and not affected by TET loss. (**a**) The presence of bacterial contaminations in wild type (*wt*) and *tet^null/+^* adult flies was checked by PCR using universal primers against bacterial 16S rDNA and against the endosymbiotic bacteria gene *wsp*. PCRs were performed on DNA from parental (F0) flies and after 3 generations of breeding in axenic conditions (F3). (**b-d**) 6mA levels were measured by LC-MS/MS in gDNA from whole larvae (b) or dissected CNS (c, d) generated from axenic flies reared on holidic medium (b,c) or conventional flies reared on classic medium (c,d). (**e**) 5mC levels were measured by LC-MS/MS in gDNA from dissected CNS from flies reared on classic medium. (**f, g**) 6mA levels were measured by LC-MS/MS in embryos (f) and dissected adult brains (g) collected from crosses with conventional or axenic (*Ax*.) individuals raised on classic fly medium supplemented (*Ax*.) or not with antibiotics. *wt:* wild type (*w^1118^*); *tet^null^*: *tet^null/null^; tet^1/2^: tet^DMAD1/DMAD2^*. Filled circles: conventional flies; open triangles: axenic flies. Individual values, means and standard deviations are plotted. No statistically significant differences were observed between *wt* and *tet* mutant samples (Mann-Whitney test).

Changes in 6mA levels following genetic manipulations of putative adenine methylases or demethylases have brought further credence to the existence and role of this modification in metazoans (Boulias & Greer, 2022). In drosophila, the absence of TET (also called DMAD, for DNA Methyl Adenine Demethylase) was associated with a strong increase in 6mA levels in embryo or adult ovary and brain (Yao et al., 2018; Zhang et al., 2015a). In addition, *in vitro* experiments suggested that drosophila TET can demethylate 6mA (Yao et al., 2018; Zhang et al., 2015a). However, that TET mediates 6mA oxidation is at odd with the well-characterized function of this family of enzymes in 5mC oxidation in metazoans (Lio *et al*., 2020). Moreover, the other enzymes involved in methyladenine oxidation/demethylation belong to the AlkB family (Boulias & Greer, 2022; Xu & Bochtler, 2020), which is related to, but distinct from the TET family (Iyer et al., 2009; Jia *et al*., 2017). Indeed, conserved residues involved in TET 5mC recognition differ from those found within the AlkB family and may not be able to accommodate a purine residue instead of a pyrimidine (Hu *et al*., 2013; Iyer et al., 2009; Parker *et al*., 2019; Xu & Bochtler, 2020). Nevertheless, the fact that the drosophila genome presents extremely low levels of 5mC and does not code for any 5mC DNA methyltransferase (Iyer *et al*., 2011; Krauss & Reuter, 2011) prompted the idea that TET could catalyze other forms of DNA modifications and notably 6mA oxidation/demethylation in the absence of its canonical substrate (Zhang et al., 2015a). As *tet* was shown to be highly expressed in the larval CNS (Delatte *et al*., 2016; Wang *et al*., 2018), we first focused on its impact on 6mA in this tissue. Yet, we found that the levels of 6mA measured by LC-MS/MS in the absence of TET (*tet^null^,* an allele that abolishes *tet* transcription (Delatte et al., 2016), were similar to wild type in the larval CNS using either axenic flies raised on holidic medium or non-axenic flies raised on classic medium (*i.e*. conventional conditions) (Fig. 1c). Of note, TET loss did not show any impact on 6mA level either in whole larvae (Fig. 1b). As previous experiments showing an increase in 6mA in the absence of TET were performed under conventional conditions with *tet^DMAD1^*/*tet^DMAD2^* mutant alleles (Yao et al., 2018; Zhang et al., 2015a), which introduce a premature stop codon in *tet* open reading frame before its catalytic domain (Zhang et al., 2015a), we repeated the analyses with this allelic combination. Yet, we did not find any increase of 6mA levels in the larval CNS in this setting either (Fig. 1d). Moreover, consistent with previous results showing that TET does not control 5mC oxidation in drosophila (Zhang et al., 2015a), we observed very low levels of 5mC (around 0.001%) in the larval CNS and no increase upon TET loss (Fig. 1e). In addition, the first product of 5mC oxidation, 5hmC (5-hydroxymethylcytosine), was below the detection limit (0.00001%). These results suggest that TET is not involved in 6mA (or 5mC) demethylation in the drosophila larval CNS.

To test whether the lack of impact of TET on 6mA levels that we observe here contrary to previous studies (Yao et al., 2018; Zhang et al., 2015a), could be due to a tissue-specific effect or to breeding conditions, we assessed 6mA levels in embryos and adult brains using conventional or germ-free flies (see Methods). As *tet^null^* homozygote mutation is pupal lethal (Zhang et al., 2015a), only *tet^DMAD1/DMAD2^* could be used for adult brains. The presence/absence of bacterial contaminants in conventional *versus* germ-free stocks was validated by PCR (Supplementary Figure S1a). Moreover, by performing gDNA sequencing, we found around 2% of bacterial DNA contaminant in gDNA in conventional flies *versus* less than 0.003% in germ-free flies (Supplementary Figure S1b). Hence, possible traces of bacterial contaminations in axenic samples should have negligible impact on LC-MS/MS measurements. LC-MS/MS analyses showed that 6mA levels were higher in embryos (Fig. 1f) or adult brains (Fig. 1g) using conventional flies as compared to their germ-free siblings. They were also more variable across samples in non-axenic conditions. It is thus likely that 6mA levels measured in non-axenic conditions do not solely reflect endogenous 6mA in the drosophila genome and variations between genotypes should be interpreted with caution. Still, we did not observe any significant increase in 6mA levels in the absence of TET in non-axenic conditions (Fig. 1f, g). Importantly, the same observation was made in axenic conditions (Fig. 1f, g). Of note, the apparent increase in 6mA levels in *tet^DMAD1/DMAD2^* axenic embryos was not reproduced in *tet^null^* embryos, suggesting that it does not simply reflect *tet* loss-of-function, and it was not statistically significant as compared to wild type levels (Welch two sample t-test: p=0.075). All together, we did not find consistent evidence that TET loss caused an increase in 6mA levels in embryos, whole larvae, larval CNS or adult brain.

As an alternate method to study TET impact on 6mA, we used SMRT sequencing (Boulias & Greer, 2022). We generated SMRT-seq data on CNS gDNA from three biological replicates of wild type and *tet^null^* larvae. As genome coverage is an important parameter to analyze SMRT-seq data, we first merged the three replicates to increase read density. In the resulting fusion datasets, around 95% of all the adenines were covered at least 25x (Supplementary Figure S2 and Table S1). The detection of 6mA by SMRT-seq is based on a modification quality value (mQV or QV), reflecting the consistency by which a specific modification is observed at a given position in a subread. Using standard parameters (coverage ≥25x and QV≥20), we found respectively, 0.158% and 0.172% of potential 6mA in the CNS of wild type and *tet^null^* larvae (Fig. 2a, Supplementary Table S2), which is much more than expected based on LC-MS/MS measurements. However, when we considered the methylation status of these adenines in the individual samples, only 2.6% were labeled as 6mA in all 3 replicates and 13.7% in at least two replicates (Fig. 2b, Supplementary Figure S3 and Table S3), consistent with the idea that SMRT-seq gives a high rate of false positives in organisms containing low 6mA levels (Douvlataniotis et al., 2020; Kong *et al*., 2023; Schadt *et al*., 2013; Zhu et al., 2018). Interestingly, increasing the QV strongly ameliorated the fraction of replication and drastically reduced the proportion of potential 6mA both in wild type and *tet^null^* datasets, whereas increasing the coverage had little effect (Fig. 2a, b, Supplementary Figure S3 and Table S3). We thus analyzed SMRT-seq results from wild type and *tet^null^* replicates using a coverage≥25x and increasing QV. However, we did not observe any significant differences either in the percentage of 6mA/A or in the fraction of methylation of these potential 6mA even with stringent QV values (Fig. 2c, d and Supplementary Table S4). Hence, while SMRT-seq data are noisy, as cautioned in previous studies, they did not reveal an increase in 6mA levels in the absence of TET.

**Figure 2.**
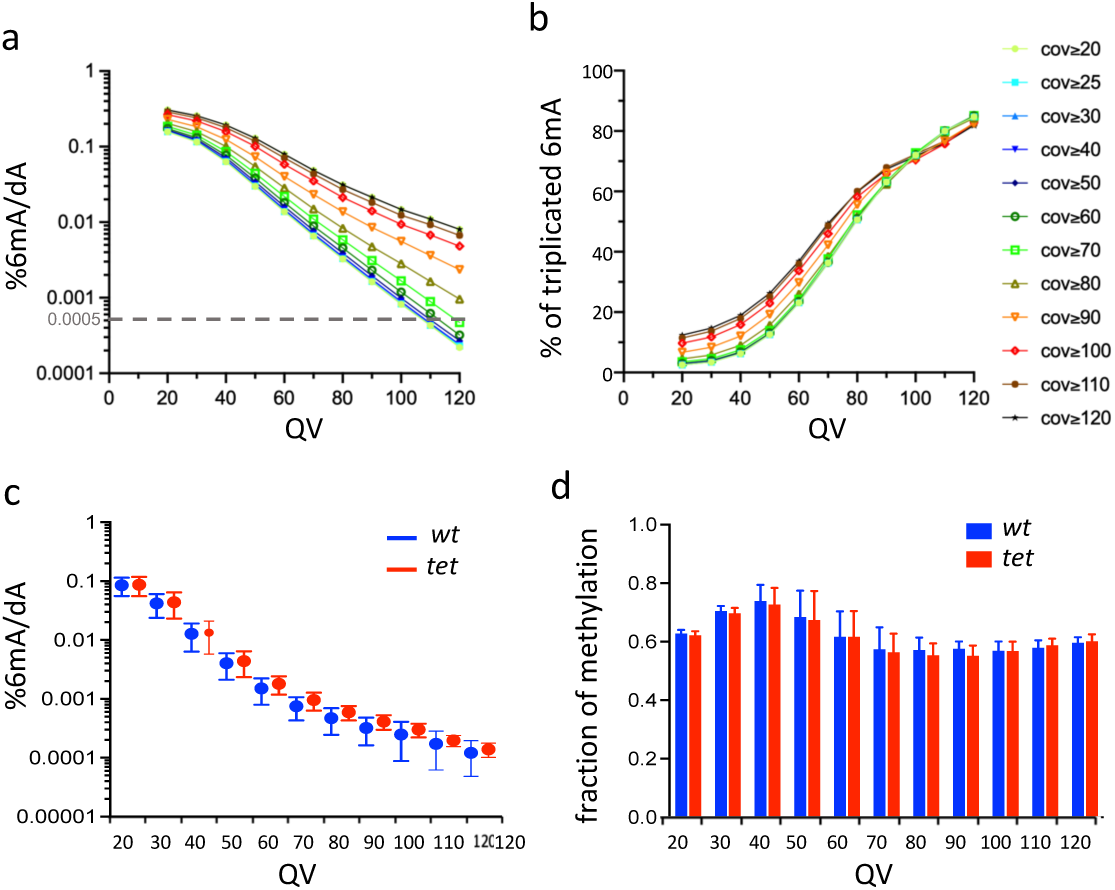
SMRT-seq analysis of larval CNS gDNA does not reveal an increase in 6mA in the absence of TET. (**a**) Percentage of adenines identified as 6mA in the wild type fusion dataset depending on the QV and coverage values used for 6mA selection. The dashed grey line indicates the level of 6mA measured by LC-MS/MS (0.0005%). (**b)** Influence of the QV and the coverage values on the proportion of 6mA identified in the wild type fusion dataset and in the three original samples. (**c**) Percentage of adenines covered at least 25x and identified as 6mA in each of the 3 wild type *(wt)* or *tet null (tet)* datasets depending on the QV. Means and standard deviations are represented. (**d**) Fraction of methylation in *wt* or *tet null* datasets depending on the QV (coverage≥25x). Means and standard deviations are represented.

To directly test whether TET demethylates 6mA, we then assessed its activity *in vitro*. Accordingly, the recombinant catalytic domain of drosophila TET was incubated with double-stranded DNA (dsDNA) containing either a 5mC or a 6mA modification and the level of modified bases was quantified by LC-MS/MS at different time points. Under our experimental conditions, 5mC levels were drastically reduced in 1 min with the concomitant appearance of 5mC oxidation products (5hmC and 5fC) (Fig. 3a). In sharp contrast, the level of 6mA remained constant even after 30 min of incubation (Fig. 3b). Hence, contrary to 5mC, 6mA does not seem to be a good substrate for drosophila TET *in vitro*. Of note, only traces of 5mC, 5hmC, 5fC or 6mA were observed when recombinant TET was incubated with non-modified dsDNA, indicating that the levels measured in the presence of modified dsDNA were not due to contaminations (Supplementary Figure S5). Our results contrast with previous reports showing that recombinant drosophila TET demethylates 6mA on dsDNA *in vitro* (Yao et al. 2018; Zhang et al., 2015a). However, both studies ran much longer reactions (up to 10 hours) and used different sources of recombinant protein (drosophila TET catalytic domain purified from human HEK293T cells). Notably, Zhang et al. (2015a) only found around 2.5% of 6mA demethylation at 30 min and less than 25% after 10 hours of incubation as measured by HPLC-MS/MS analyses. These results suggest that drosophila TET may oxidize 6mA, but with a much lower affinity than 5mC since with observed a near complete oxidation of 5mC after 1 min. and no significant decrease in 6mA levels after 30 min. of reaction (for identical concentrations of substrate and enzyme). It is possible too that the preparation of TET catalytic domain in different systems changes its enzymatic activity, potentially in relation to distinct post-translational modifications. Still, it is worth mentioning that a distant TET homolog in the fungus *Coprinopsis cinerea* was recently shown to oxidize both 5mC and 6mA (Mu *et al*., 2022). Importantly, its peculiar capacity to bind and demethylate 6mA requires key residues within its catalytic domain which are not conserved in other TET homologs including in drosophila (Supplementary Figure S6). These observations support our conclusion that drosophila TET does not serve as 6mA demethylase.

**Figure 3.**
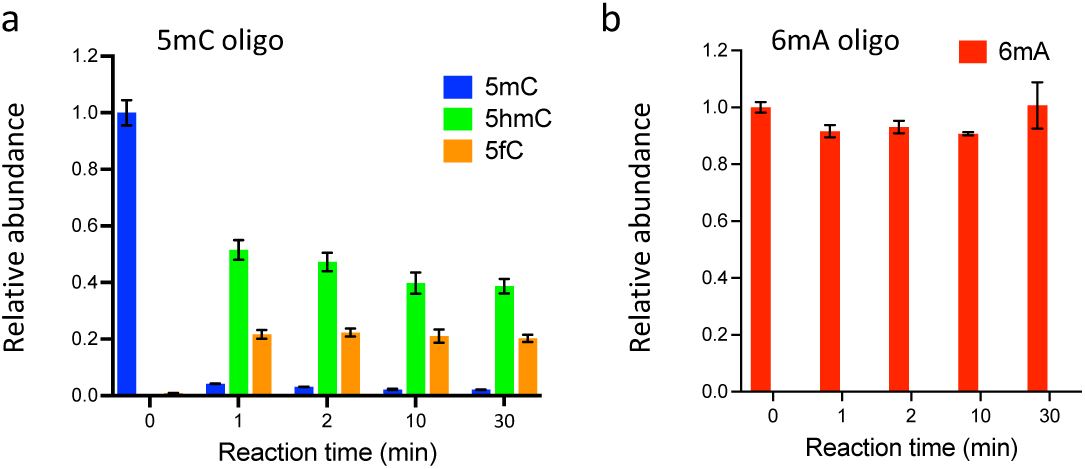
TET does not oxidize 6mA. (**a**, **b**) *In vitro* assays showing TET activity profile on 5mC (a) and 6mA (b) containing double stranded oligonucleotide substrates. The levels of 5mC and its oxidised products (5hmC and 5fC) are represented relative to 5mC level at t=0. The levels of 6mA are represented relative to 6mA level at t=0. Error bars denote standard deviations from 3 independent experiments.

Although previous reports suggested that TET controls fly viability, ovarian development or adult brain formation by demethylating 6mA (Yao et al., 2018; Zhang et al., 2015a), the functional importance of TET enzymatic activity has never been tested genetically as available *tet* alleles either abolish its expression or delete the whole catalytic domain. In view of our results and to address this issue, we generated a catalytic dead mutant allele of *tet* (*tet^CD^*). Accordingly, the conserved HxD iron-binding motif required for the catalytic activity of TET/AlkB dioxygenase family of enzymes (Hu et al., 2013; Tahiliani *et al*., 2009; Xu & Bochtler, 2020) was mutated by CRISPR/Cas9-mediated homologous recombination using a *tet-GFP* knock-in which allows to tag all protein TET isoforms with GFP (Fig. 4a, b). Of note, immunostaining in the larval CNS confirmed that TET is widely expressed in this tissue and showed that the H1947Y/D1949A mutation does not alter TET expression or its nuclear localization (Fig. 4c-h). Strikingly, while *tet^null^* homozygote individuals die at the pupal stage (Delatte et al., 2016), we found that *tet^CD/CD^* as well as *tet^CD/null^* pupae had a normal hatching rate and gave rise to viable adult flies (Fig. 4i). We did not observe lethality of *tet^CD/CD^*individuals at earlier developmental stage either (Supplementary Figure S7). Then, we assessed whether this mutation affected adult wing positioning, ovarian development or mushroom body formation as reported upon TET loss of expression (Wang et al., 2018; Yao et al., 2018; Zhang et al., 2015a). Yet, *tet^CD/CD^* flies did not exhibit the “held out” wing positioning defects present in *tet^DMAD1/DMAD2^* adult escapers (Fig. 4j, j’, j’’). Similarly, atrophied ovaries or mushroom body projection defects observed in the absence of TET expression were not reproduced when only its catalytic activity was impaired (Fig. 4k, k’, k’’ and 4l, l’, l’’). Thus, TET function in drosophila seems essentially independent of its enzymatic activity, indicating that TET-mediated regulation of 6mA level, if it truly happens, is not essential either for fly development.

**Figure 4.**
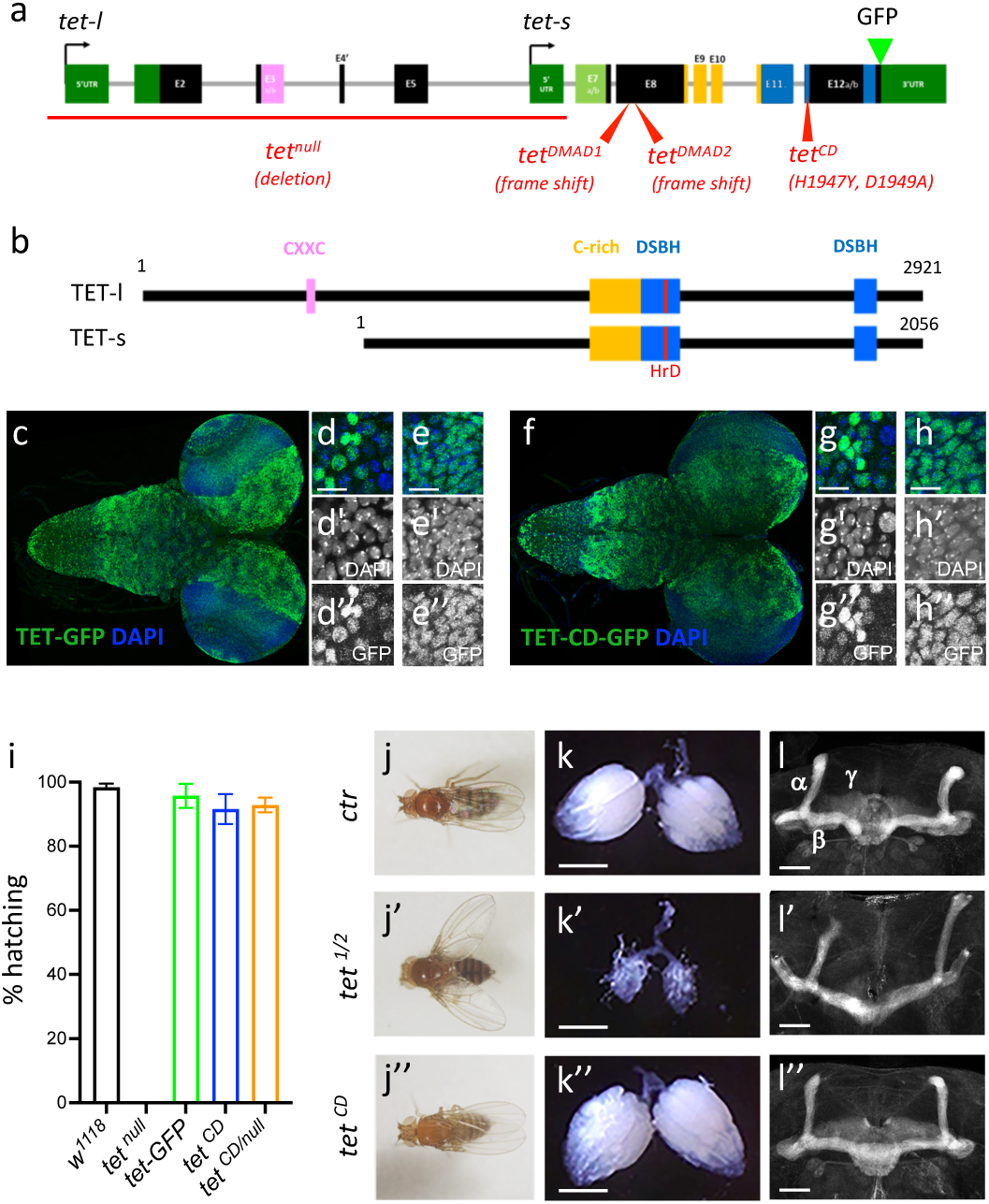
TET catalytic activity is largely dispensable in drosophila. (**a, b**) Schematic representation of *tet* locus (a) and main protein isoforms (b). (**a**) *tet* is transcribed from two alternative promoters giving rise to *tet-long (tet-l)* and *tet-short (tet-s*) isoforms. Filled boxes represent exons; non-coding exons (UTR) are depicted in green, and coding exons in black or according to their domain-associated color. Introns are represented as grey lines (not to scale). The *tet null, DMAD1, DMAD2* and *catalytic dead (CD*) alleles are depicted in red. The location of the GFP insertion generated by CRISRP/Cas9-mediated knock-in is also indicated. (**b**) The conserved domains of TET are colored; pink: CXXC DNA binding domain, orange: Cystein-rich domain, blue: double-stranded ß helix (DSBH) domain, red: HxD (iron binding motif). Amino acid positions are indicated according to the longest TET-l and TET-s isoforms. (**c-h**) Expression pattern of the wild type and catalytic-dead versions of TET in the larval CNS. *tet-GFP* (c-e) and *tet^CD^-GFP* (f-h) knock-in lines were used to detect TET proteins by confocal imaging after immunostaining against GFP (green). Nuclei were labelled with DAPI (blue). (c, f): stitched images showing dorsal views of the entire CNS. Scale bar: 100 µm. (d, e, g, h): high-magnification views of TET expression in the ventral nerve cord (d, g) or the central brain (e, h). DAPI-only and GFP-only channels are presented in the middle (‘) and lower (‘’) panels, respectively. Scale bar: 10 µm. (**i**) Percentage of adult flies of the indicated genotypes hatching from their pupal case. Means and standard deviations from 4 independent experiments. (**j-l**) Wing positioning (j), ovaries (k) and mushroom bodies (l) of wild type adult flies (*ctr: tet-GFP)* as compared to flies lacking TET expression (*tet^1/2^*: *tet^DMAD1/DMAD2^* adult escapers) or TET enzymatic activity (*tet^CD^*). (l-l”) Immunostaining against Fas2 on adult brains was used to label mushroom body a, b and g lobes. (k-k”) Scale bar 500µm. (l-l”) Scale bar 50µm.

## Conclusions

In sum, our results confirm that 6mA is present only at very low levels in the drosophila genome. With only a few hundred methylated adenines per haplogenome, we argue that 6mA is unlikely to play a major regulatory function in normal conditions. In addition, we did not find any evidence that TET loss promotes 6mA accumulation. Rather, our results strongly suggest that this conserved enzyme is not a methyladenine demethylase and that its catalytic activity is largely dispensable for drosophila development. Further experiments will be necessary to firmly establish whether regulated adenine methylation/demethylation takes place and what are the enzymes involved, not only in drosophila but also in other metazoans. Besides, our analyses call for further investigations of the molecular mechanisms underlying the essential, enzymatic-independent, mode of action of TET.

## Methods

### Fly strains and breeding

The following *D. melanogaste*r strains were used: *w^1118^*(control, Bloomington), *tet^null^* (Delatte et al., 2016), *tet^DMAD1^*, *tet^DMAD2^* (Zhang *et al*., 2015a). *tet^null^* (a kind gift from Dr R. Steward) was generated by FRT site recombination between two PBac insertions, resulting in the absence of *tet* transcription (Delatte et al., 2016). *tet^DMAD1^* and *tet^DMAD2^* (kind gifts from Dr D. Chen) were obtained by CRISPR and produce truncated TET proteins deleted from their C-terminal domain (including the whole catalytic domain) (Zhang *et al*., 2015a). The *tet-GFP* knock-in line was generated by InDroso Functional Genomics (Rennes, France) using CRISPR/Cas9-mediated homologous recombination to insert the EGFP in frame with the last amino acid of TET. Similarly, the catalytic dead *tet^CD^* flies were generated by CRISPR/Cas9-mediated homologous recombination in the *tet-GFP* background to mutate TET HRD motif into YRA (*dm6:* chromosome 3L:2,791,624 “A” to “C” and 3L:2,791,631 “C” to “T”). In both cases, the resulting flies were validated by sequencing.

Unless otherwise specified, stock maintenance and sample collection were performed using classic fly medium (75 g/l organic corn flour, 28 g/l dry yeast, 40 g/l sucrose, 8 g/l agar, 10 ml/l Moldex 20%) with a 12 h dark:light cycle. Germ-free drosophila lines were generated as described (Sabat *et al*., 2015). Briefly: embryos were collected on grape juice agar plates, dechorionated with 2.7% bleach for 2-3 min, washed in sterile ddH2O and transferred to standard fly medium supplemented with antibiotics (50 µg/ml amoxicillin, tetracyclin, kanamycin and puromycin) for at least two successive generations. When “holidic” medium was used to avoid any source of contamination by exogenous DNA, embryos from germ-free adults were collected on grape juice agar plates, dechorionated, washed with ddH2O and transferred to a chemically-defined medium, using the amino acid ratio of the FLYAA recipe (Piper *et al*., 2017), together with antibiotics to maintain axenic conditions. All crosses and larvae collections were performed at 25°C.

### Viability assays

Embryos were collected at 25°C from 1-week-old flies over 8h on grape juice agar plates. For each genotype, batches of 100 embryos were transferred to corn flour-yeast-agar plates; the number of first instar larvae was counted after 30h, the number of pupae was counted at day 9 and the number of hatched adults was counted from days 10 to 15. Each experiment was repeated at least four times.

### Immunostainings

Third instar larvae or adult fly brains were dissected in 1X Phosphate Buffer Saline (PBS) and fixed for 25 min in PBS containing 4% paraformaldehyde. Fixed samples were washed rapidly twice with PBS and 3 times for 15 min with PBS-0.3% Triton X-100 (PBT) before being pre-incubated for 1h in PBT-1% bovine serum Albumin (BSA, Sigma). Samples were incubated overnight at 4°C with primary antibody diluted in PBT-1% BSA, washed 3 times for 15 min in PBT, and incubated with respective secondary antibodies diluted in PBT-1% BSA for 3h at room temperature or overnight at 4°C. Samples were washed in PBT and mounted in Vectashield-DAPI (Vector Laboratories). Images were acquired using a Leica LSM800 confocal microscope. The following antibodies were used: goat anti-GFP (Abcam, 1/500), mouse anti-Fasciclin II (DSHB, 1/25), donkey anti-goat Alexa fluor 488 (Invitrogen, 1/1000), donkey anti-mouse Cy3 (Jackson Immuno, 1/1000).

### Bacterial contamination assays

The presence of bacterial DNA contamination in parental and “germ-free” derived stocks was checked by PCR using universal primers targeting bacterial *16S rDNA* (*16S-s*: 5’-AGAGTTTGATCCTGGCTCAG-3’, *16S-r:* 5’-GGTTACCTTGTTACGACTT-3’ (Weisburg *et al*., 1991)) and primers amplifying the *wsp* gene from the endosymbiont *Wolbachia* (*wsp-s*: 5’-TGGTCCAATAAGTGATGAAGAAAC-3’, *wsp-r*: 5’-AAAAATTAAACGCTACTCCA-3’ (as from (Jeyaprakash & Hoy, 2000)). The presence of any contaminant was also checked by DNA sequencing: for each sample, genomic DNA from 10 adult flies (5 males/5 females) was extracted using DNeasy Blood & Tissue Kit (Qiagen). DNA was resuspended and sheared in 1X TE (0.1 mM EDTA, 10 mM Tris HCl pH 8.0) by sonication for 20 min (30 sec ON/30 sec OFF) using the Bioruptor Pico (Diagenode) to obtain an average size of ∼300pb. DNA libraries were prepared from 1µg DNA using the NEBNext Ultra II DNA Library Prep Kit (Illumina) following manufacturer’s instructions and sequencing was performed by Novogene (Cambridge, UK) using NovaSeq 6000 (paired-end, 150pb). Between 15 to 19 million reads were obtained per sample. To assess the presence of contamination, the resulting reads were first aligned to the drosophila reference genome (dm6 Ensembl release 70) with Bowtie2. Unaligned reads were then processed for blast search to bacteria, viral and fungal genomes using the DecontaMiner tool (Sangiovanni *et al*., 2019).

### LC-MS/MS analyses

For DNA purification, whole larvae, bleach-dechorionated embryos or dissected adult brains of the required genotypes were washed in sterile PBS, crushed in lysis buffer (100 mM Tris-HCl pH 9, 100 mM EDTA, 1% SDS), incubated at 70°C for 30 min and then in 1 M potassium acetate at 4°C for 30 min. After centrifugation at 12000 g for 20 min, the supernatant was digested with RNAse A and RNAse H for 3 h at 37°C, extracted twice with phenol-chloroform-isoamyl alcohol (25:24:1) and precipitated with isopropanol. DNA from dissected third instar larval brains (around 100 per sample) was extracted using DNeasy Blood & Tissue Kit (Qiagen), digested with RNAse A and RNAse H for 3 h. In both types of extraction, DNA was precipitated with 500 mM ammonium acetate (pH 6.2). DNA pellets were dissolved in ddH2O, and their concentration, as the absence of RNA contamination, were checked on a Qubit 3.0 fluorometer (Invitrogen).

Up to 1200 ng of DNA per sample were digested to nucleosides using 0.6 U nuclease P1 from *P. citrinum* (Sigma-Aldrich), 0.2 U snake venom phosphodiesterase from C*. adamanteus* (Worthington), 0.2 U bovine intestine phosphatase (Sigma-Aldrich), 10 U benzonase (Sigma-Aldrich), 200 ng Pentostatin (Sigma-Aldrich) and 500 ng Tetrahydrouridine (Merck-Millipore) in 5 mM Tris (pH 8) and 1 mM MgCl2 for 2 h at 37°C. 1000 ng of digested DNA were spiked with internal standard (D_3_-5mC and D_2_,^15^N_2_-5hmC, 250 fmol each) and subjected to analysis by LC-MS (Agilent 1260 Infinity system in combination with an Agilent 6470 Triple Quadrupole mass spectrometer equipped with an electrospray ion source (ESI)). The solvents consisted of 5 mM ammonium acetate buffer (pH 5.3, adjusted with acetic acid; solvent A) and LC-MS grade acetonitrile (solvent B; Honeywell). A C18 reverse HPLC column (SynergiTM 4 µM particle size, 80 Å pore size, 250 × 2.0 mm; Phenomenex) was used at a temperature of 35°C and a constant flow rate of 0.5 mL/min was applied. The compounds were eluted with a linear gradient of 0-20% solvent B over 10 min. Initial conditions were regenerated with 100% solvent A for 5 min. The four main nucleosides were detected photometrically at 254 nm *via* a diode array detector (DAD). The following ESI parameters were used: gas temperature 300°C, gas flow 7 L/min, nebulizer pressure 60 psi, sheath gas temperature 400°C, sheath gas flow 12 L/min, capillary voltage 3000 V, nozzle voltage 0 V. The MS was operated in the positive ion mode using Agilent MassHunter software in the MRM (multiple reaction monitoring) mode. Therefore, the following mass transitions were used to detect the respective modifications: 6mA 266->150; 5mC 242->126; D_3_-5mC 245->129; 5hmC 258->142; D_2_,^15^N_2_-5hmC 262->146; 5fC 256->140. For absolute quantification, internal and external calibrations were applied as described previously (Kellner *et al*., 2014), except for 6mA and 5fC, for which only external calibration was performed.

### SMRT sequencing analyses

Around 250 brains from *w^1118^* or *tet^null^*third instar larvae were dissected for each sample. Tissues were crushed in lysis buffer (100 mM Tris-HCl pH 9, 100 mM EDTA, 100 mM NaCl,1% SDS), digested with RNase A for 15 min at room temperature and 30 min at 65°C, before being incubated with 4 volumes of 3.2 M LiCl, 0.9 M KAc for 30 min at 4°C. After centrifugation at 12000 g for 20 min at 4°C, the supernatant was extracted twice with phenol-chloroform-isoamyl alcohol (25:24:1) and DNA was precipitated with isopropanol. 5 µg of DNA were used to prepare each sequencing library.

SMRT-seq was performed at the Gentyane Sequencing Platform (Clermont-Ferrand, France) with a PacBio Sequel Sequencer (Pacific Biosciences, Menlo Park, CA, USA). The SMRTBell libraries were prepared using a SMRTbell Express 2 Template prep kit, following the manufacturer’s recommendations. High Molecular Weight Genomic DNA (5 µg) was sheared with the 40 kb program using a Diagenode Megaruptor (Diagenode) to generate DNA fragments of approximately 30 kb. Assessment of the fragment size distribution was performed with a Femto Pulse (Agilent Technologies, Santa Clara, CA, USA). Sheared genomic DNA was carried into enzymatic reactions to remove single-strand overhangs and to repair any damage that may be present on the DNA backbone. An A-tailing reaction followed by the overhang adapter ligation was conducted to generate the SMRTBell templates. After a 0.45X AMPure PB beads purification, the samples were size-selected using the BluePippin (Sage Science, Beverly, MA, USA) to recover all the material above 15 kb. The samples were then purified with 0.45X AMPure PB Beads to obtain the final libraries of around 30 kb. The SMRTBell libraries were quality inspected and quantified on a Femto Pulse and a Qubit fluorimeter with Qubit dsDNA HS reagent Assay kit (Life Technologies). A ready-to-sequence SMRTBell Polymerase Complex was created using a Binding Kit 3.0 (PacBio) and the primer V4, the diffusion loading protocol was used, according to the manufacturer’s instructions. The PacBio Sequel instrument was programmed to load a 6 pM library and samples were sequenced on PacBio SMRTCells v2.0 (Pacific Biosciences), acquiring one movie of 600 min per SMRTcell.

For each sequenced sample, SMRT-seq reads were aligned using pbmm2 tool (https://github.com/PacificBiosciences/pbmm2) on Drosophila genome (dm6 Ensembl release 70). For each condition (wild type or *tet^null^*), a fusion of alignments of the three biological replicates was done using « samtools merge » (Li *et al*., 2009). 6mA detection was performed on individual and on merged samples with IpdSummary tool from KineticTools (http://github.com/PacificBiosciences/kineticsTools) applying the following parameters: --identify m6A --numWorkers 16 --pvalue 0.01 –identifyMinCov 5 –methylFraction. To detect 6mA with higher confidence, we applied several thresholds on coverage and modificationQV (QV) with homemade scripts in bash and R. The SMRT-seq data are deposited under GEO accession number GSE206852.

### Purification of drosophila TET catalytic domain

The catalytic domain of *drosophila* TET (dTET) was cloned in pET28a expression vector. The His-tagged protein was overexpressed in *E. coli* BL21 (DE3) CodonPlus RIL cells for 17 h at 16°C. Cells were resuspended in lysis buffer (50 mM HEPES pH 7.5, 20 mM imidazole, 500 mM NaCl, 1 mM DTT, 10% glycerol and supplemented with protease inhibitor 0.2 mM PMSF) and disrupted using Bandelin Sonoplus ultrasonic homogenizer. The cell lysates were cleared by centrifugation (Lynx 600 (Thermo), Fiberlite F21-8×50y) at 38.300 g for 30 min at 4°C and the supernatant was loaded onto an affinity column packed with Ni-NTA agarose beads (Genaxxon, Germany). The column was washed with wash buffer (50 mM HEPES pH 7.5, 20 mM imidazole, 500 mM NaCl, 1 mM DTT, 10% glycerol) then the recombinant protein was eluted with elution buffer (50 mM HEPES pH 7.5, 250 mM imidazole, 500 mM NaCl, 1 mM DTT, 10% glycerol). Purified protein was dialyzed against dialysis buffer I (50 mM HEPES pH 7.5, 1 mM DTT, 300 mM NaCl and 10% glycerol) followed by dialysis buffer II (50 mM HEPES pH 7.5, 1 mM DTT, 300 mM NaCl and 50% glycerol).

### TET activity assays

The double-stranded DNA substrates were prepared by annealing forward oligos and reverse complement counterpart by heating at 95°C followed by bringing temperature to RT slowly on the heat block in Annealing Buffer (10mM Tris-HCl pH7.5 100mM NaCl). Forward oligos sequences containing a 5mC or 6mA modified nucleotide were the following: 5mC 5’-GTAAGTCTGGCA5mCGTGAGCCTCAGAG-3’, 6mA 5’-GTAAGTCTGGCG6mAGTGAGCCTCAGAG-3’. The reaction was performed with 0.5 μM DNA substrate and 2 μM recombinant dTET in Reaction Buffer (50 mM HEPES pH 6.8, 100 μM Ammonium ion(II) sulfate hexahydrate, 1 mM, α-ketoglutarate, 1 mM ascorbic acid and 150 mM NaCl) at 37 °C. Reaction was stopped at different time points by adding 2µl of 0.5 M EDTA to a 40µl volume reaction followed by heating at 90°C for 5 min. Samples were treated with proteinase K for 1 h at 50°C and precipitated with ethanol. The level of 6mA, 5mC, 5hmC and 5fC was analyzed by LC-MS/MS as described above.

## Authors’ contribution

LV and LW conceived the study. MB, GG, YR, LV and LW designed the experiments. MB, GG, EP, MSD, RB and XN performed the experiments. MB, GG, YR, MSD, TJ, MH, LV and LW analyzed the results. LW and LV wrote the manuscript with inputs from all co-authors. All authors read and approved the final manuscript.

## Acknowledgements

We are grateful to Dr. A. Molaro for critical reading of the manuscript and to the members of the iGReD for helpful discussions. We also thank Dr V. Gautier and the Gentyane platform at the Clermont-Ferrand INRAE for PacBio sequencing and the iGReD CLIC imaging facility for help with confocal experiments. We thank the Drosophila Bloomington Stock Center, Dr. R. Steward (Piscataway), Dr. D. Chen (Beijing) for fly stocks, as well as the Developmental Study Hybridoma Bank for antibodies. This project was supported by grants from the Agence Nationale de la Recherche (ANR-17-CE12-0030-03), Fondation ARC (PJA20171206371) and I-Site Cap20-25 (EpiMob) to LW. MB was supported by fellowships from the Université Clermont-Auvergne and the Fondation pour la Recherche Médicale (FRM). GG was supported by fellowships from the Université Clermont-Auvergne and the Ligue Nationale Contre le Cancer. MH was funded by the Deutsche Forschungsgemeinschaft (DFG, German Research Foundation) TRR-319 TP C03, SPP1784, HE 3397/13-2 and HE 3397/14-2.

---

**Supplementary Figure S1.**
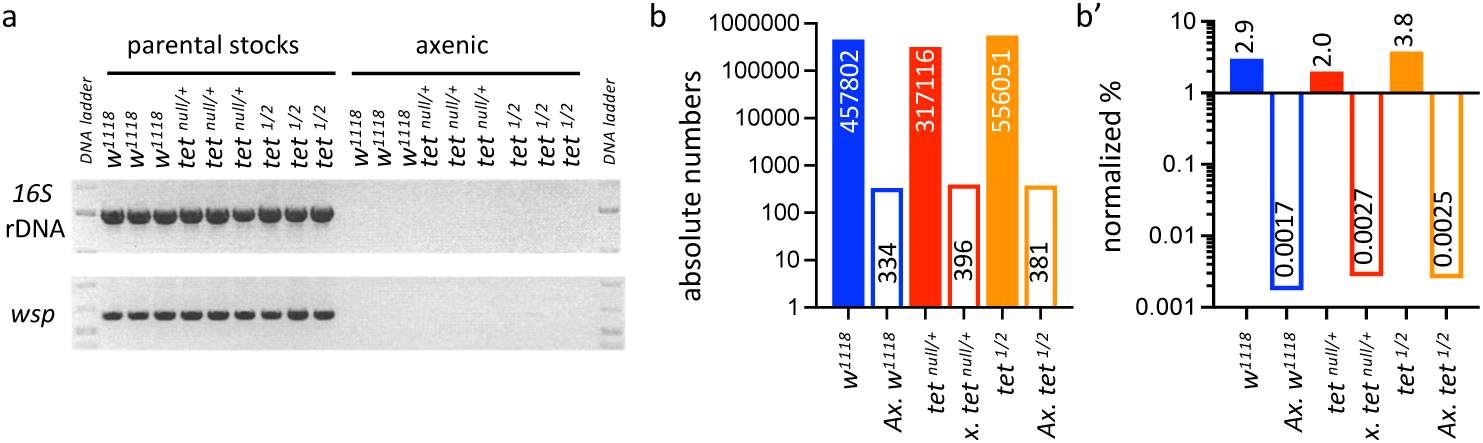
Detection of bacterial contamination in parental stocks and after 3 generations of breeding in axenic conditions. (**a**) The presence of bacterial contamination in adult flies of the indicated genotypes was checked by PCR using universal primers against bacterial *16S rDNA* and against the endosymbiotic bacteria gene *wsp*. (**b, b’**) Genome-wide sequencing was used to assess the presence of contamination in gDNA from adult flies of the indicated genotypes. Between 15 and 19 million reads were analysed per sample. The absolute numbers of reads mapping to bacteria, virus or fungi genomes are represented in b, and their proportions normalized to the total number of reads are presented in b’. *tet^1/2^: tet^DMAD1/DMAD2^*.

**Supplementary Figure S2.**
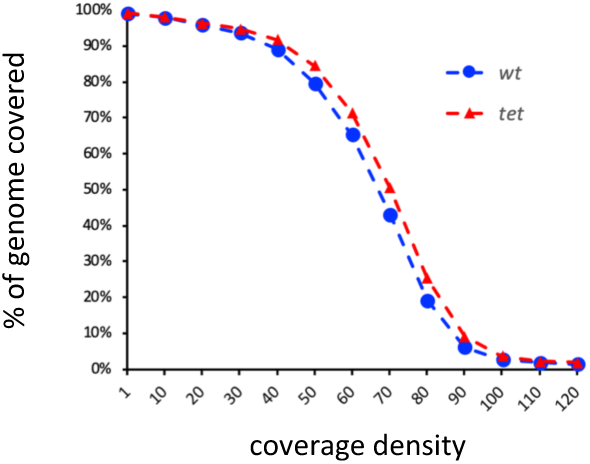
Proportion of the drosophila genome covered in *wild type* (*wt*) and *tet^null^* (*tet*) SMRT-seq fusion datasets according to coverage density.

**Supplementary Figure S3.**
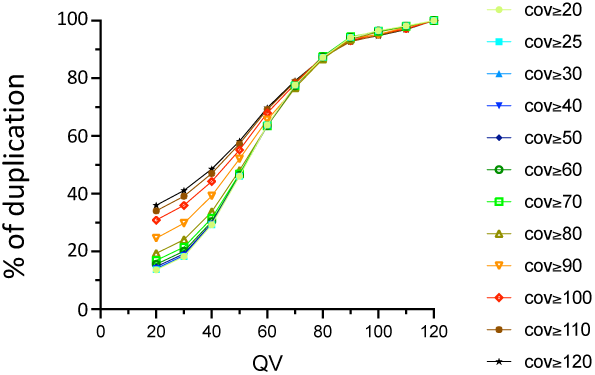
Influence of the QV and the coverage values on the proportion of 6mA identified in the *wt* fusion dataset and in the three original samples. The percentage of 6mA identified in at least 2 out of 3 replicates is shown.

**Supplementary Figure S4.**
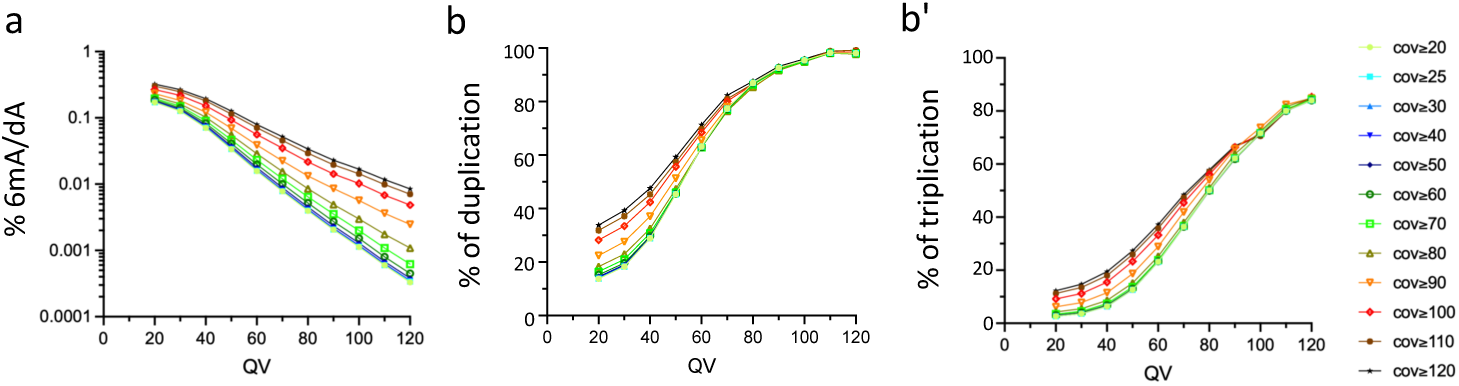
(**a**) Percentage of all adenines identified as 6mA by SMRT-seq in the *tet^null^* fusion dataset depending on the QV and coverage values used for 6mA selection (log10 scale). (**b, b’**) Influence of the QV and the coverage values on the proportion of 6mA identified in the *tet^null^* fusion dataset and in the three original samples (b: percentage of 6mA identified in at least 2 out of 3 samples; b’: percentage of 6mA identified in all 3 samples.

**Supplementary Figure S5.**
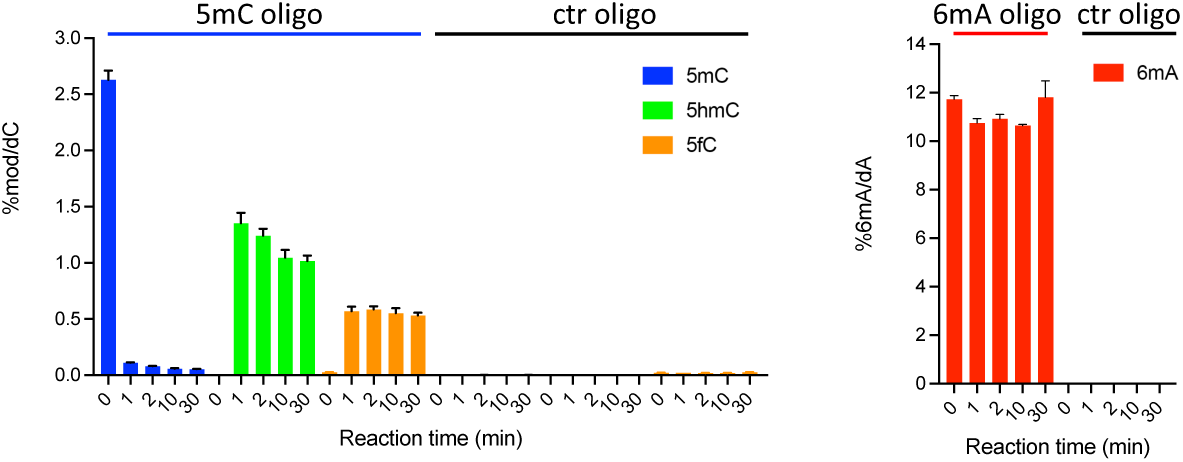
*In vitro* assays showing TET activity profile on double stranded oligonucleotide substrates containing or not 5mC (**a**) or 6mA (**b**). The levels of 5mC, 5hmC and 5fC normalized to dC (a) or 6mA normalized to dA (b) are represented. Error bars denote standard deviations from 3 independent experiments. Only background levels of modified nucleosides were detected when purified recombinant TET catalytic domain was incubated with unmodified (ctr) oligonucleotides.

**Supplementary Figure S6.**
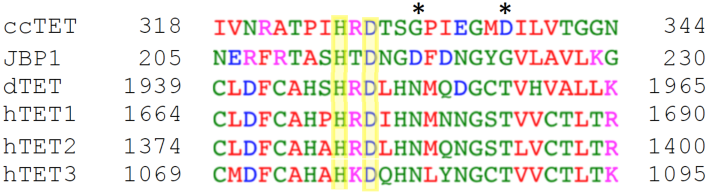
Multiple sequence alignment of TET/JBP family members. The sequence surrounding the HxD motif in *Coprinopsis cinerea* TET (ccTET), *Trypanosoma brucei* JBP1, *Drosophila melanogaster* TET (dTET) and *Homo sapiens* TET1, TET2, TET3 is shown. Conserved amino-acids between TET homologs are boxed in yellow. The two key amino acids required for 6mA oxidation by ccTET are labelled with a star.

**Supplementary Figure S7.**
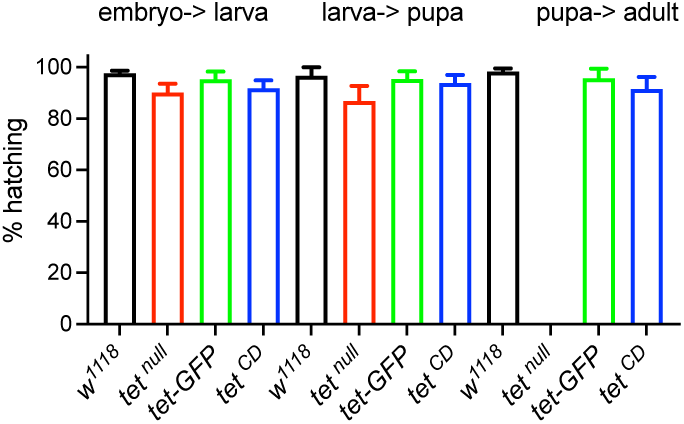
Survival assays showing the percentage of hatching embryos, larvae and adults of the indicated genotypes. Means and standard deviations are from at least 6 independent experiments.

